# Quality control of low-frequency variants in SARS-CoV-2 genomes

**DOI:** 10.1101/2020.04.26.062422

**Authors:** Mikhail Rayko, Aleksey Komissarov

## Abstract

During the current outbreak of COVID-19, research labs around the globe submit sequences of the local SARS-CoV-2 genomes to the GISAID database to provide a comprehensive analysis of the variability and spread of the virus during the outbreak. We explored the variations in the submitted genomes and found a significant number of variants that can be seen only in one submission (singletons). While it is not completely clear whether these variants are erroneous or not, these variants show lower transition/transversion ratio. These singleton variants may influence the estimations of the viral mutation rate and tree topology. We suggest that genomes with multiple singletons even marked as high-covered should be considered with caution. We also provide a simple script for checking variant frequency against the database before submission.

## Introduction

Sequencing of viral genomes allowed researchers to track the distribution of the viruses on Earth, and to assess the rate of the viral evolution. This task is especially important during the active outbreaks, where arisen mutations may affect test systems and vaccines under development.

During the current pandemic of SARS-nCoV-2, the primary resource for consolidating genomic data is the GISAID database (Shu et al., 2017). As a result of the collaborative efforts of the researchers worldwide, on April 14, 2020 it contained over 8,000 SARS-nCoV-2 genomes from different countries, sequenced and assembled using various technologies and approaches.

Unfortunately, these sequences are not error-free. Different sequencing technologies are characterized by different types and frequency of errors (Ma et al., 2019). Often these sequencing errors are not random and are typical for certain sets of nucleotides such as homopolymers. At least 21% of submitted genomes on April 1,2020 are sequenced using Oxford Nanopore technology according to GISAID (according to searching by “nanopore” or “minion” keywords in metadata). Oxford Nanopore technology is error-prone and ONT data requires careful polishing.

Another source of systematic errors can be the use of a fixed set of primers, which leads to the enrichment of some regions over others. While many assemblers imply more or less uniform coverage, primer sets (such as described at https://github.com/CDCgov/SARS-CoV-2_Sequencing) are commonly used to enrich the sequences. The use of PCR enrichment may result in low coverage of individual genomic regions (even in case of high average coverage), which can be a serious source of errors in the downstream analysis.

In addition to sequencing errors, the variety of genome assembly methods makes it difficult to compare data, and the lack of access to raw data makes it impossible to reassemble data using a standardized approach. Prompt access to original raw sequencing data is needed to perform accurate and reproducible analysis.

GISAID database curators do a tremendous job of filtering submitted sequences, but sometimes it is difficult to distinguish real variants from errors, especially at the lack of information about coverage. Here we compared variants across the submissions and developed a pipeline to separate real variants from potential errors based on their frequency across all genomes in the database. We suppose that variations observed in a single genome from the dataset - hereinafter referred to as *singletons* - may be erroneous, and one should proceed with caution, or maybe even filter out singleton-containing genomes from downstream applications until we get additional evidence from other samples.

## Methods

### Dataset

8,053 full-length (>29,000 bp) sequences of the SARS-CoV-2 were downloaded from the GISAID database (www.epicov.org) on April 14, 2020, including 5,556 genomes marked as “high coverage”.

### Variation calling

Sequences were aligned to the reference genome (NCBI RefSeq NC_045512.2) using minimap2 (Li, 2018). Resulting vcf files were merged using MergeVcfs tool from Picard toolkit (http://broadinstitute.github.io/picard/). SNVs were annotated using SnpEff 4.3t (Cingolani et al.,2012). Ts/Tv was calculated as a direct transition/transversion ratio on a filtered set of SNVs (not considering their multiplicity, and excluding indels and Ns). Scripts for data analysis and visualization are available at https://github.com/ablab/covid19_variation_analysis.

### Sequencing and assembly technology statistics

We used the following keywords for GISAID database to get information about sequencing methods: “Illumina”, “Nanopore”, “Ion Torrent”, “Sanger”, “dbnseq”. We used the following keywords for GISAID database to get information about assembly methods: “artic”, “phe”, “spades”, “dnbseq”, “megahit”, “clc”, “ivar”, and “seattle”. To get various assembly methods based on raw reads mapping we used the following keywords: “mpileup”, “bwa”, “bowtie”, or “mapping”.

## Results

### Submitted sequences contain a large fraction of singleton variations

After filtering out all variants containing Ns, there are variants in 4,562 positions (out of 29,903 bp). 3,006 of them were identified as singletons. Figure 1 illustrates quantity and distribution over the genome for SNVs of different multiplicity.

**Figure 1.**
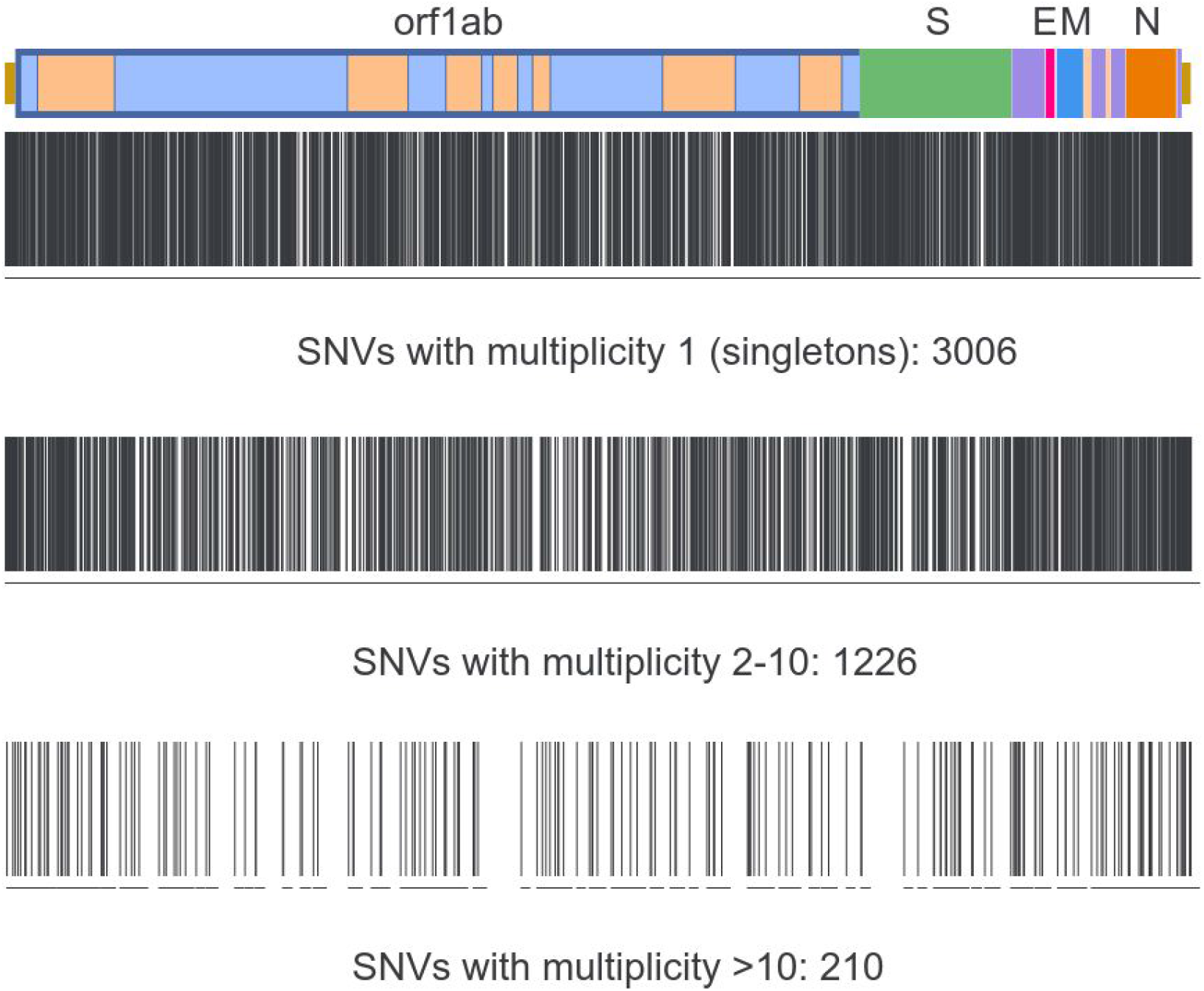
Visualisation of the obtained SNVs in SARS-CoV-2 genomes collected before April 14, 2020. Top to bottom: singletons, SNVs observed in 2-10 genomes, SNVs observed more than in 10 genomes.

### Singleton variations show decreased Ts/Tv ratio

We explored transition/transversion (Ts/Tv) ratio for the variants observed with different frequencies. For singletons this ratio is lower than for more frequent variants. (Table 1). Lower Ts/Tv ratio corresponds to false positive results (e.g. Wang et al. 2015, Guo et al 2012), and may indicate the introduced sequencing/assembly errors.

**Table 1.** Ts/Tv for variants that occur in SARS-nCoV-2 genomes with different frequencies.

| Variant Frequency | All genomes |  |  | “High coverage” |  |  |
| --- | --- | --- | --- | --- | --- | --- |
|  | Ts | Tv | Ts/Tv | Ts | Tv | Ts/Tv |
| 1 (singleton) | 1,834 | 1,132 | 1.62 | 1,355 | 697 | 1.94 |
| 2 | 492 | 214 | 2.3 | 364 | 141 | 2.58 |
| 3 | 214 | 97 | 2.21 | 175 | 75 | 2.33 |
| 4 | 125 | 47 | 2.66 | 92 | 32 | 2.88 |
| 5 | 80 | 25 | 3.2 | 44 | 15 | 2.93 |
| >=6 | 311 | 114 | 2.73 | 247 | 92 | 2.68 |

Currently, genomes with more than 0.05% singleton mutations (i.e. more than 15 SNPs) are automatically excluded from “high-covered” in GISAID database. When comparing the fraction of genomes marked as “high-covered” among genomes that contain singletons we see that for genomes with 2 and less singletons there is no significant difference. However, the fraction of genomes with 3 or more singletons is significantly (p<0.01, counted with χ2 criteria) lower than in those that do not contain any singletons (see Table 2).

**Table 2.**
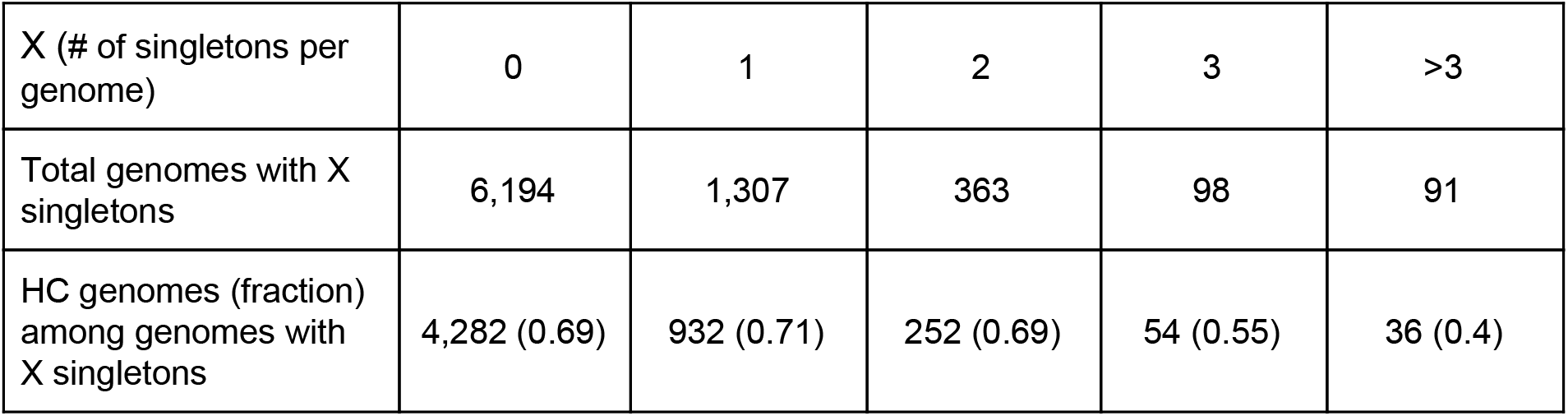
Number of genomes with different amounts of singletons, and their fraction in genomes marked as “high-covered” in GISAID database (denoted as “HC”).

| X (# of singletons per genome) | 0 | 1 | 2 | 3 | >3 |
| --- | --- | --- | --- | --- | --- |
| Total genomes with X singletons | 6,194 | 1,307 | 363 | 98 | 91 |
| HC genomes (fraction) among genomes with X singletons | 4,282 (0.69) | 932 (0.71) | 252 (0.69) | 54 (0.55) | 36 (0.4) |

**Table 2.** Types of mutations in the coding regions.

|  | Singletons | Non-singletons | Singletons from HC genomes | Non-singletons from HC genomes |
| --- | --- | --- | --- | --- |
| In-frame indels | 42 (1.44% ) | 15 (0.9%) | 2 (0.1%) | 5 (0.3%) |
| Frameshift indels | 101 (3.5%) | 5 (0.3%) | 9 (0.5%) | 3 (0.2%) |
| Missense variants | 1,838 (62.8%) | 970 (59.5%) | 1,064 (63.7%) | 859 (58.9%) |
| Synonymous variants | 891 (30.5%) | 630 (38.6%) | 575 (34.4%) | 582 (39.9%) |
| Stop gained mutations | 52 (1.8%) | 11 (0.7%) | 18 (1%) | 9 (0.62%) |
| Stop lost mutations | 1 (0.1%) | 0 | 1 (0.1%) | 0 |
| Total mutations in coding regions | 2,925 | 1,631 | 1,669 | 1,458 |

### Frameshift indels and nonsense mutations are usually correspond to singletons

All variants in coding regions were annotated with SnpEff. Singletons showed slightly higher presence of the frameshift indels and stop-gained mutations, most probably erroneous (see Table 3). Also we see a significant difference in the percentage of synonymous variants between singleton and non-singleton SNPs. However, it is not clear whether this difference corresponds to errors or not.

**Table 3.**
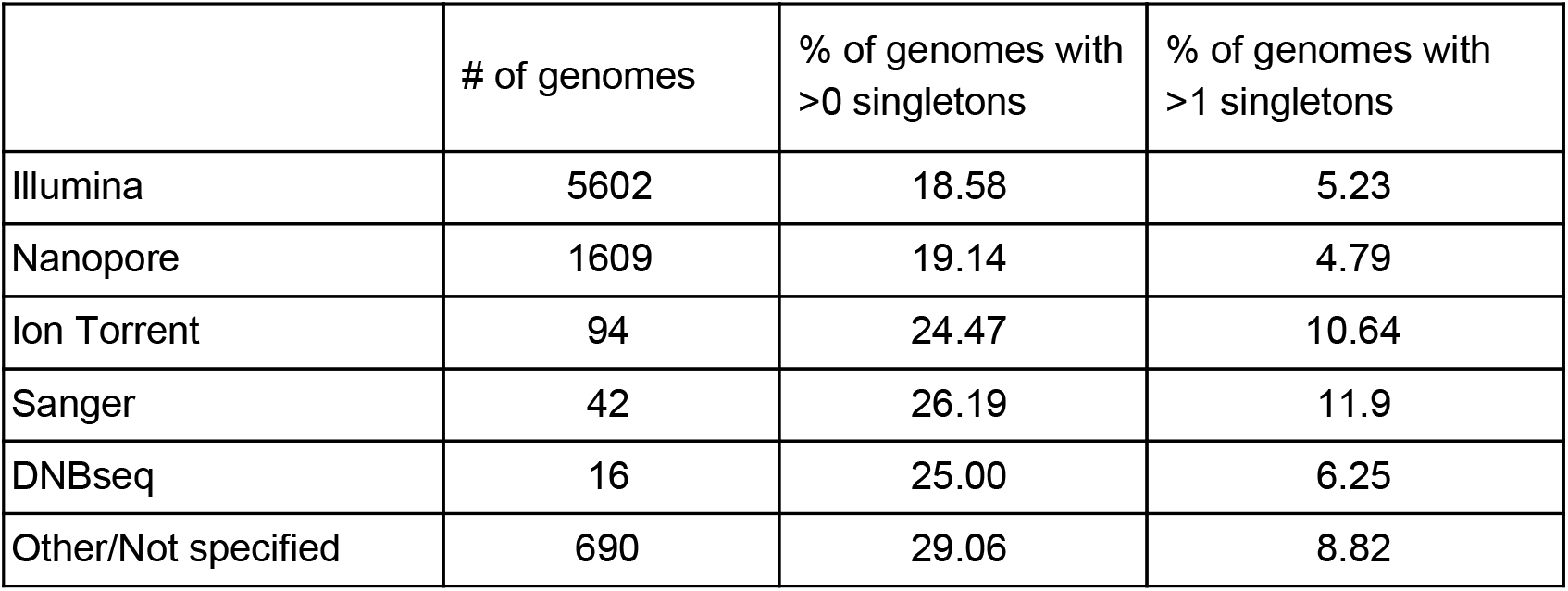
Percentage of singleton-containing genomes depending on sequencing technology.

### Indels should be verified prior to submission

Genomes marked as “high covered” in the GISAID database must not contain indels unless verified by the submitter. Thus, this is important to compare obtained indels with those already presented in the database. Out of 227 indels from samples collected and submitted to GISAID before April 14st, 2020, we observed only 33 non-singletons (see Supplemental Table 1). 30 of these non-singleton indels are observed in at least one HC genome, some of them were already described and checked (Bal et al., 2020, Su et al., 2020).

### Genome assembly method seems more important than sequencing technology

We extracted information about sequencing technology by keywords in metadata, and estimated the number of genomes with singletons. We were expecting to see an elevated number of singleton-containing genomes in the Oxford Nanopore results. However, it turned out that the proportion of the singleton-containing genomes for Illumina and ONT data is almost the same (see Table 3). There is a small amount of Sanger and DNBseq data in the database at the moment, these results may change over time.

Then we compared different assembly methods (see Table 4). We found the lowest number of singletons in genomes assembled by specialized virus-tailored pipelines, such as Artic Network (https://artic.network/ncov-2019), Phe (Public Health England), iVAR (Grubaugh et al., 2019) and Seattle flu assembly pipeline (https://github.com/seattleflu/assembly). De novo assembly with MEGAHIT (Li et al., 2015) shows a significant amount of singletons - one should probably interpret such results with caution. The full data shown in Supplementary table 2.

**Table 4.**
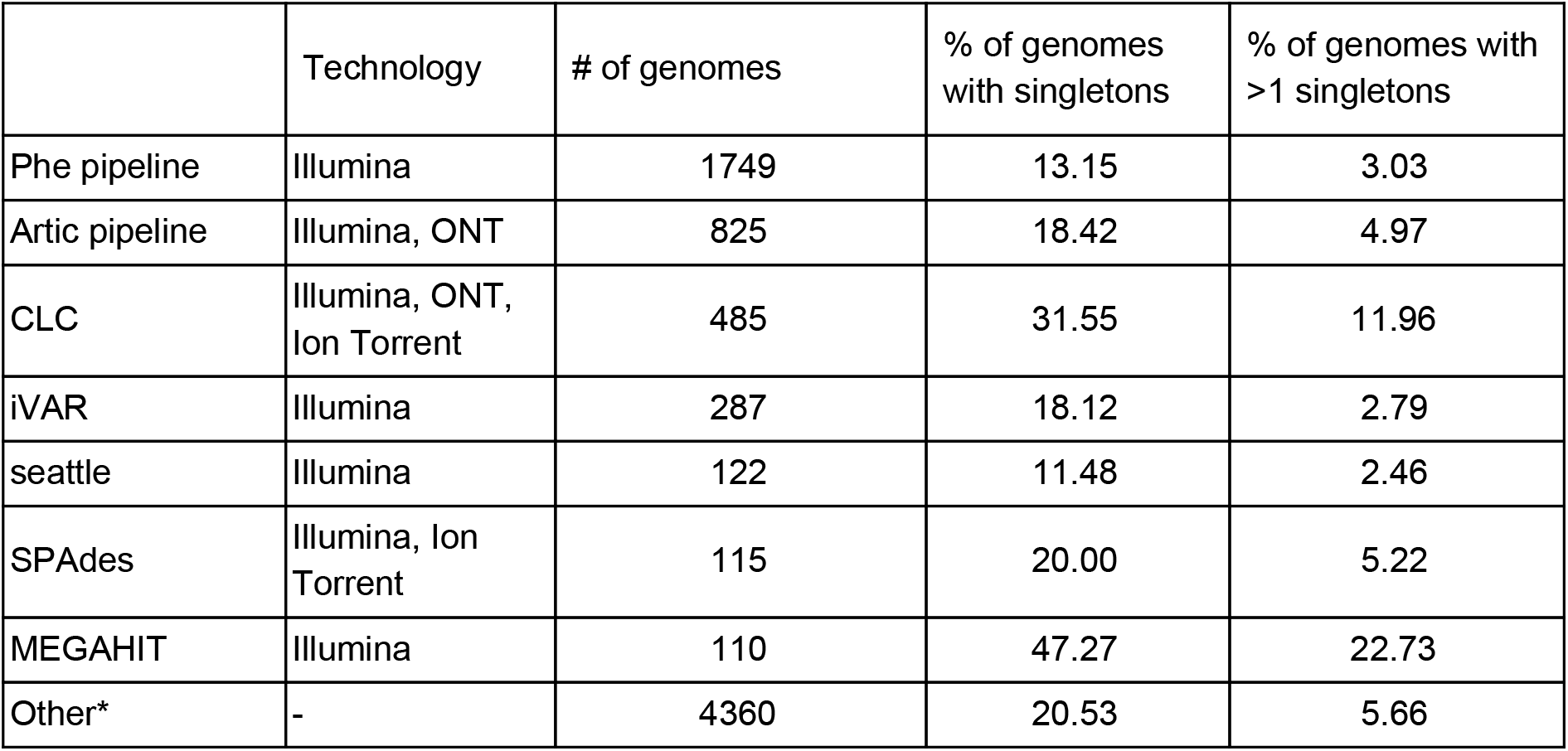
Percentage of singleton-containing genomes depending on assembly method. *“Other” category includes all custom pipelines, rarely used tools and samples with incomplete or absent information about assembly methods.

|  | Technology | # of genomes | % of genomes with singletons | % of genomes with >1 singletons |
| --- | --- | --- | --- | --- |
| Phe pipeline | Illumina | 1749 | 13.15 | 3.03 |
| Artic pipeline | Illumina, ONT | 825 | 18.42 | 4.97 |
| CLC | Illumina, ONT, Ion Torrent | 485 | 31.55 | 11.96 |
| iVAR | Illumina | 287 | 18.12 | 2.79 |
| seattle | Illumina | 122 | 11.48 | 2.46 |
| SPAdes | Illumina, Ion Torrent | 115 | 20.00 | 5.22 |
| MEGAHIT | Illumina | 110 | 47.27 | 22.73 |
| Other* | - | 4360 | 20.53 | 5.66 |

### There is a two week delay between sample collection and genome availability

For each day from January 1 to April 15 we computed a number of total known variants and a number of variants shared in more than one genome to this date (Figure 1). We found that the data on the new variants has a two week delay. This time delay should also be taken into account when analyzing the data, especially if one links it to other more rapidly updated data, such as infection statistics.

**Figure 2.**
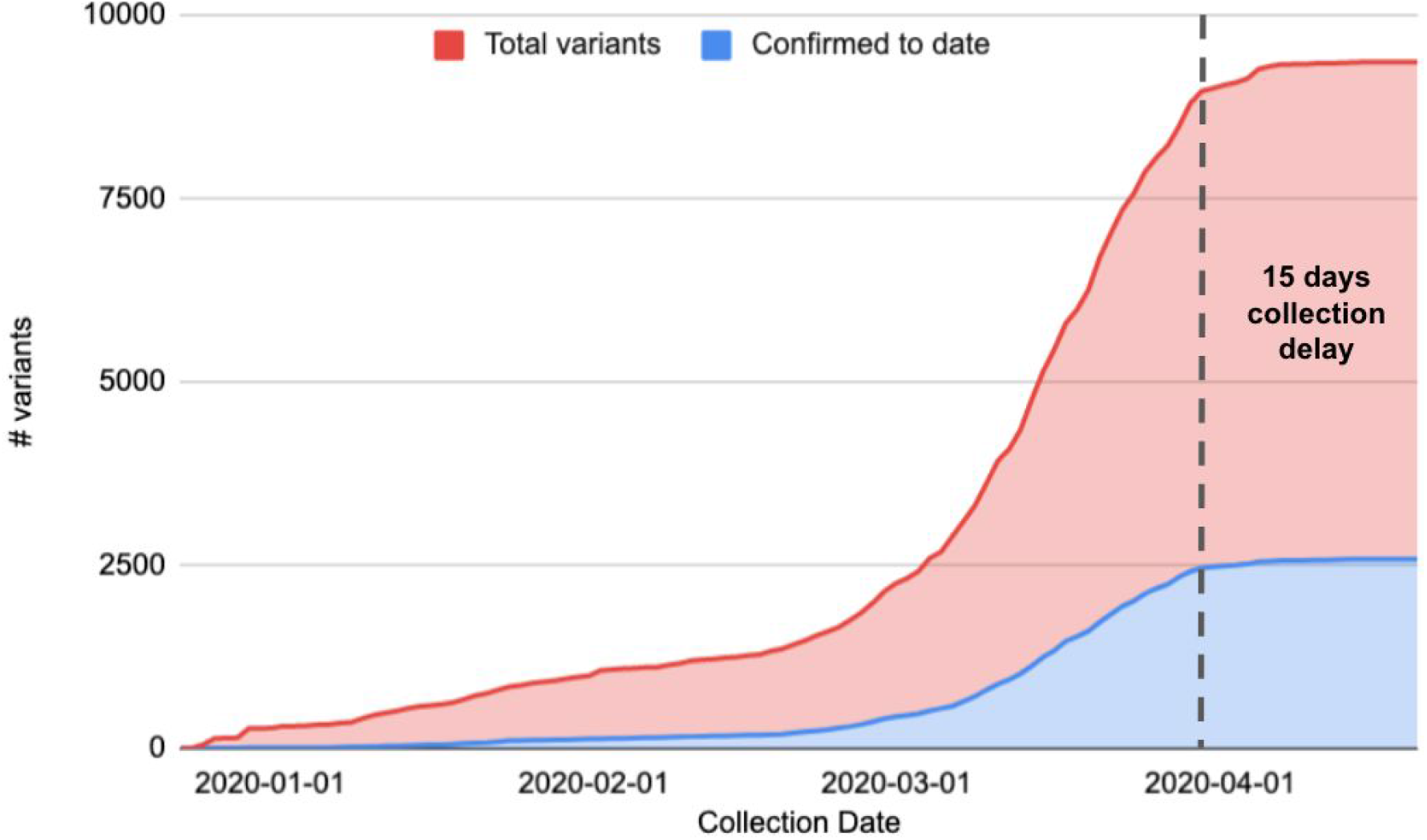
Fraction of confirmed variants during the four months period.

## Discussion

Based on our results, we suggest that the singletons can serve as indicators of the potential erroneous assembly. We provide a script that quickly allows to obtain sample’s SNVs frequencies against the current SARS-CoV-2 samples database and to use this result as a sanity check before submission. It is definitely possible to see in a recently sequenced sample a real SNV that is not present in the database yet. However, if you see plenty of them we recommend checking these locations (i.e. in some genome visualization software like Tablet (Milne et al., 2012)).

Described artifacts may influence even the estimations of the viral mutation rate and phylogenetic tree topology. We suggest that genomes with multiple singletons even marked as high-covered should be used with caution.

Although described methods allow to notice some potential errors in the database, reliable quality control for individual samples is not possible without access to reads. We hope to extend this work when more raw sequencing data connected to GISAID genomes will become available in public databases.

## Data availability

Scripts for reproducing the analysis steps and checking variant frequency against the database before submission are available at https://github.com/ablab/covid19_variation_analysis.

## Acknowledgements

Our results acknowledge, as the original source of the data, the laboratories where the clinical specimens and/or virus isolates were obtained (see full list in the <u>Supplementary table 3</u>). Aleksey Komissarov was financially supported by the Government of the Russian Federation through the ITMO Fellowship and Professorship Program. Mikhail Rayko was supported by St. Petersburg State University (ID 51555639).

We are grateful to Dmitry Antipov, Sonya Garushyants, Dmitry Meleshko, Anton Korobeynikov and Alla Lapidus for suggestions and comments that improved the paper.

---

**Supplemental Table 1.**
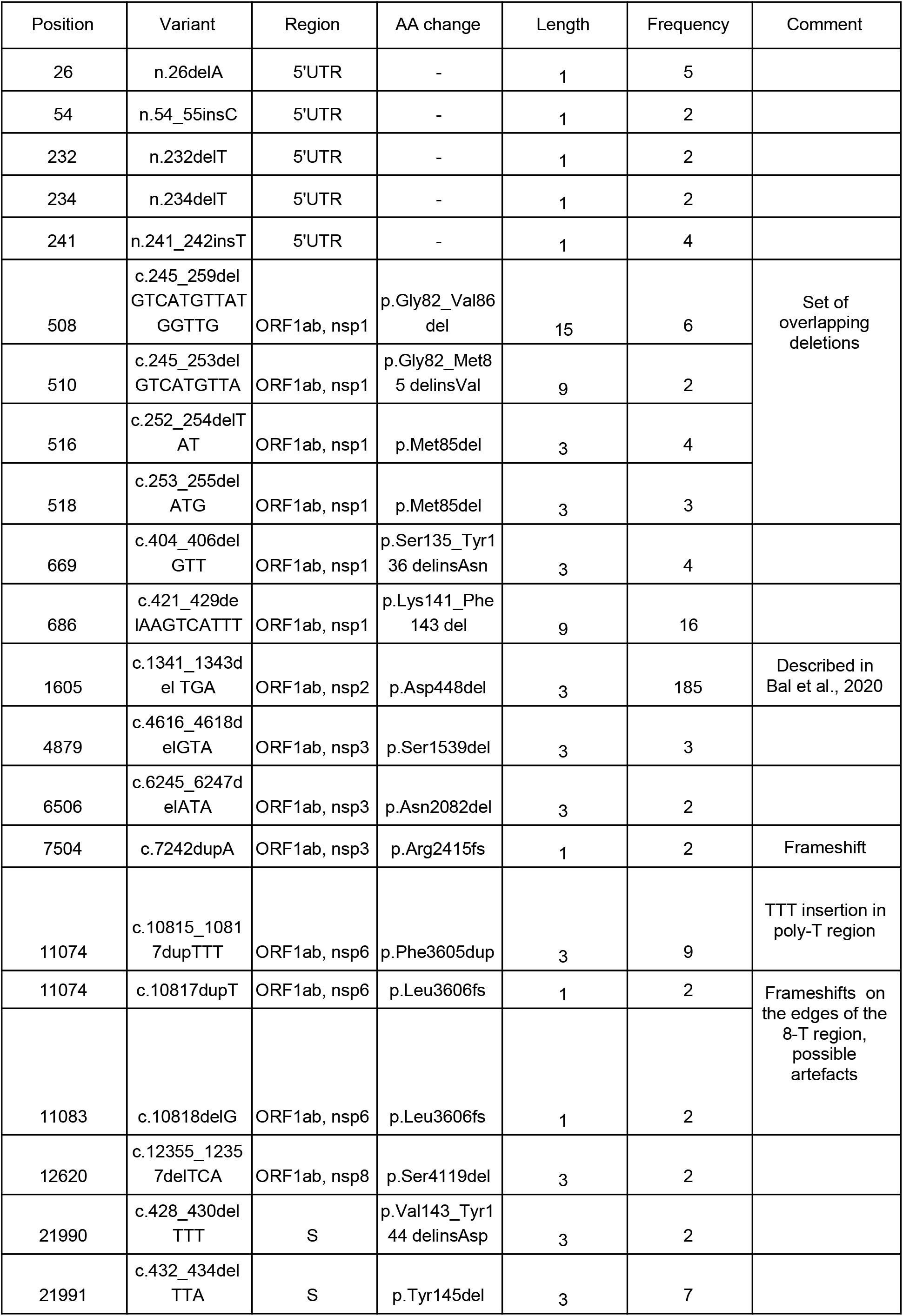
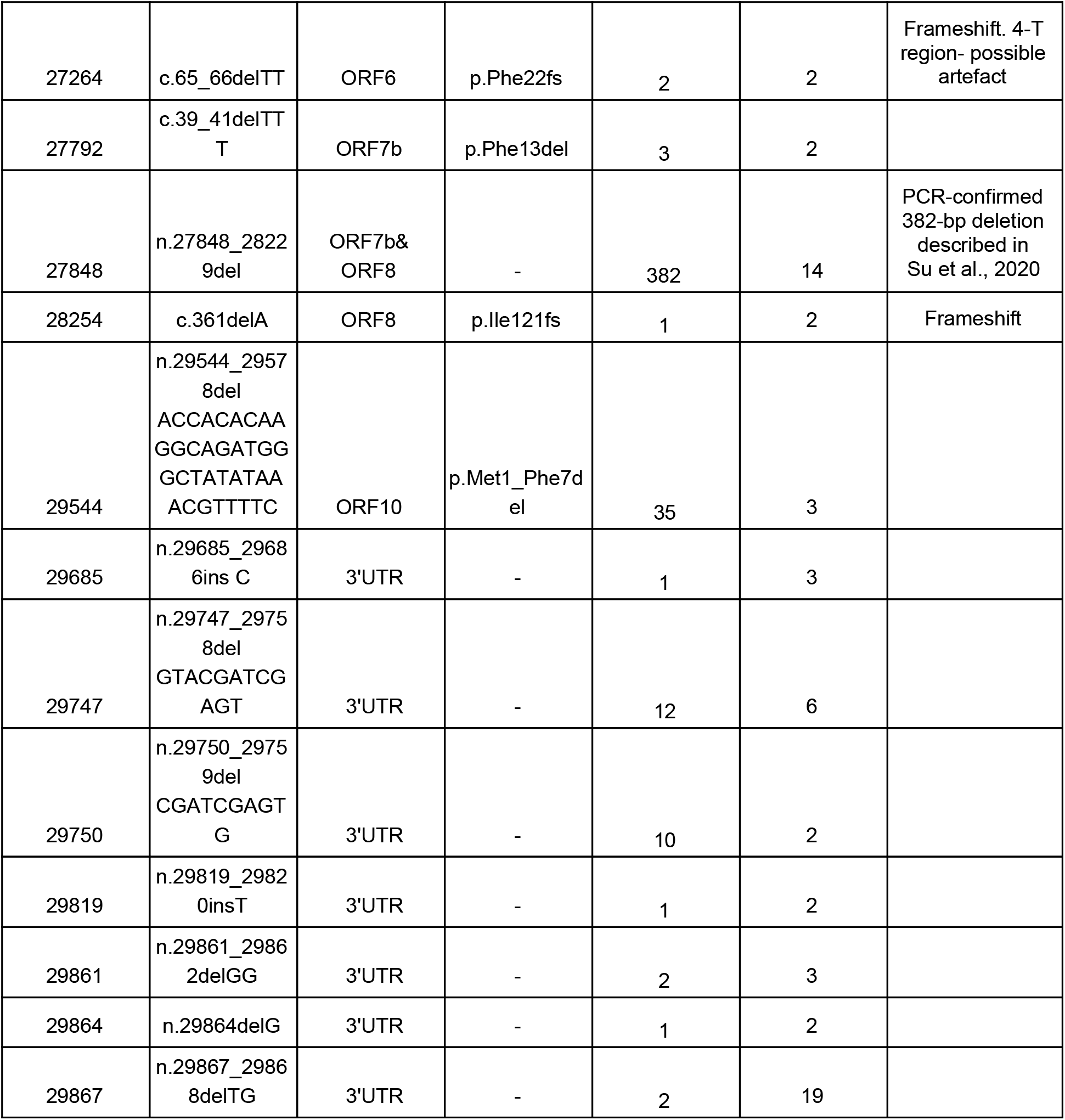
Non-singleton indels in SARS-CoV-2 genomes from GISAID database. The most frequently observed indel is the Asp448del in ORF1ab, nsp2 peptide, described by Bal et al., 2020 as Asp268Del 1607-1609. But, according to our analysis, there is a deletion between two codons, AATGAC, resulting in replacing Asn(AAT)Asp(GAC) to Asn(AAC). Another notable variant is a 382-bp deletion g.27848_28229del, almost completely removing ORF8, described by Su et al., 2020, supported by 14 samples from Singapore. Totally there are 4 indels that can be seen in 10 or more samples, 8 indels in 5 or more samples, 17 indels in 3 or more samples and 16 indels confirmed by just two samples. Some of these indels may affect protein functions - such as p.Val143_Tyr144delinsAsp and p.Tyr145del in spike protein and deletion of the first 7 amino acids in ORF10.

**Supplemental Table 2.**
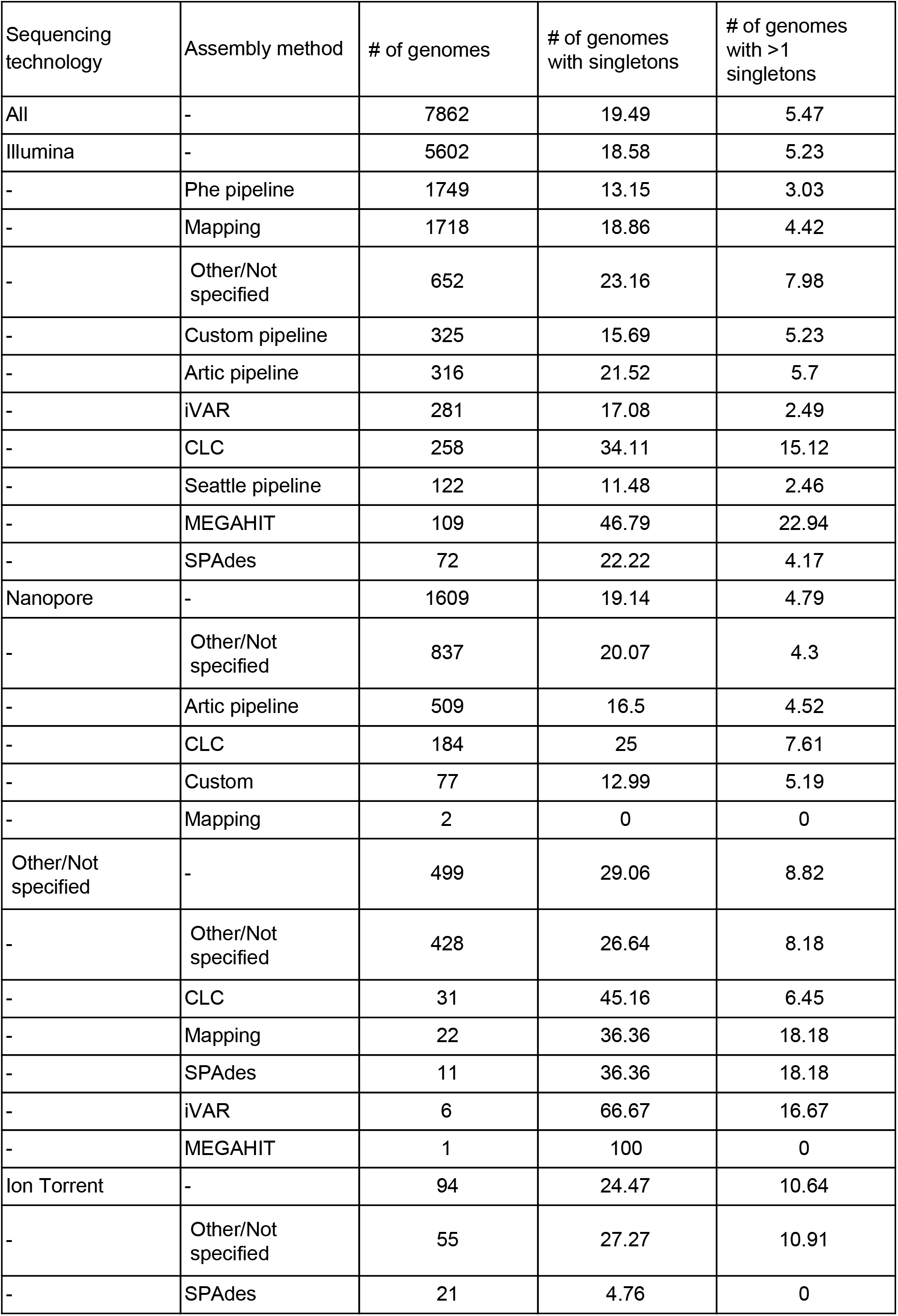
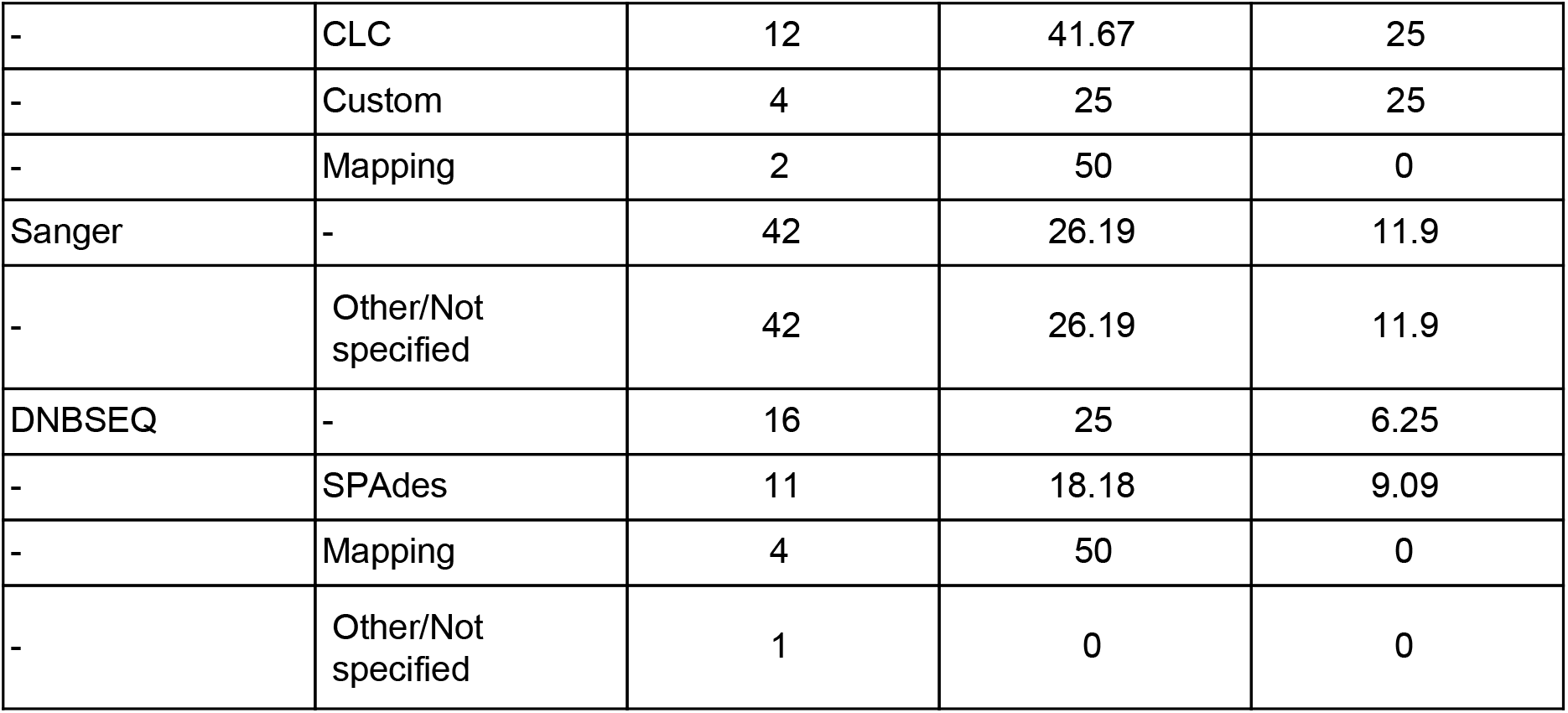
Full table with percentage of singleton-containing genomes depending on sequencing and assembly method. “Other” category includes all custom pipelines, rarely used tools and samples with incomplete or absent information about assembly methods. “Mapping” category includes all assembly methods with words “mpileup”, “bwa”, “bowtie”, or “mapping” in the description.

