## Supplemental Table 1 for "Quality control of low-frequency variants in SARS-CoV-2 genomes"

| Position | Variant | Region | AA change | Length | Frequency | Comment |
|---|---|---|---|---|---|---|
| 26 | n.26delA | 5'UTR | - | 1 | 5 | |
| 54 | n.54_55insC | 5'UTR | - | 1 | 2 | |
| 232 | n.232delT | 5'UTR | - | 1 | 2 | |
| 234 | n.234delT | 5'UTR | - | 1 | 2 | |
| 241 | n.241_242insT | 5'UTR | - | 1 | 4 | |
| 508 | c.245_259del GTCATGTTAT GGTTG | ORF1ab, nsp1 | p.Gly82_Val86 del | 15 | 6 | Set of overlapping deletions |
| 510 | c.245_253del GTCATGTTA | ORF1ab, nsp1 | p.Gly82_Met8 5 delinsVal | 9 | 2 | |
| 516 | c.252_254delT AT | ORF1ab, nsp1 | p.Met85del | 3 | 4 | |
| 518 | c.253_255del ATG | ORF1ab, nsp1 | p.Met85del | 3 | 3 | |
| 669 | c.404_406del GTT | ORF1ab, nsp1 | p.Ser135_Tyr1 36 delinsAsn | 3 | 4 | |
| 686 | c.421_429de lAAGTCATTT | ORF1ab, nsp1 | p.Lys141_Phe 143 del | 9 | 16 | |
| 1605 | c.1341_1343d el TGA | ORF1ab, nsp2 | p.Asp448del | 3 | 185 | Described in Bal et al., 2020 |
| 4879 | c.4616_4618d elGTA | ORF1ab, nsp3 | p.Ser1539del | 3 | 3 | |
| 6506 | c.6245_6247d elATA | ORF1ab, nsp3 | p.Asn2082del | 3 | 2 | |
| 7504 | c.7242dupA | ORF1ab, nsp3 | p.Arg2415fs | 1 | 2 | Frameshift |
| 11074 | c.10815_1081 7dupTTT | ORF1ab, nsp6 | p.Phe3605dup | 3 | 9 | TTT insertion in poly-T region |
| 11074 | c.10817dupT | ORF1ab, nsp6 | p.Leu3606fs | 1 | 2 | Frameshifts on the edges of the 8-T region, possible artefacts |
| 11083 | c.10818delG | ORF1ab, nsp6 | p.Leu3606fs | 1 | 2 | |
| 12620 | c.12355_1235 7delTCA | ORF1ab, nsp8 | p.Ser4119del | 3 | 2 | |
| 21990 | c.428_430del TTT | S | p.Val143_Tyr1 44 delinsAsp | 3 | 2 | |

| | | | | | | |
|---|---|---|---|---|---|---|
| 21991 | c.432_434del TTA | S | p.Tyr145del | 3 | 7 | |
| 27264 | c.65_66delTT | ORF6 | p.Phe22fs | 2 | 2 | Frameshift. 4-T region- possible artefact |
| 27792 | c.39_41delTT T | ORF7b | p.Phe13del | 3 | 2 | |
| 27848 | n.27848_2822 9del | ORF7b& ORF8 | - | 382 | 14 | PCR-confirm ed 382-bp deletion described in Su et al., 2020 |
| 28254 | c.361delA | ORF8 | p.Ile121fs | 1 | 2 | Frameshift |
| 29544 | n.29544_2957 8del ACCACACAA GGCAGATGG GCTATATAA ACGTTTTC | ORF10 | p.Met1_Phe7d el | 35 | 3 | |
| 29685 | n.29685_2968 6ins C | 3'UTR | - | 1 | 3 | |
| 29747 | n.29747_2975 8del GTACGATCG AGT | 3'UTR | - | 12 | 6 | |
| 29750 | n.29750_2975 9del CGATCGAGT G | 3'UTR | - | 10 | 2 | |
| 29819 | n.29819_2982 0insT | 3'UTR | - | 1 | 2 | |
| 29861 | n.29861_2986 2delGG | 3'UTR | - | 2 | 3 | |
| 29864 | n.29864delG | 3'UTR | - | 1 | 2 | |
| 29867 | n.29867_2986 8delTG | 3'UTR | - | 2 | 19 | |

**Supplemental Table 1.** Non-singleton indels in SARS-CoV-2 genomes from GISAID database. The most frequently observed indel is the Asp448del in ORF1ab, nsp2 peptide, described by Bal et al., 2020 as Asp268Del 1607-1609. But, according to our analysis, there is a deletion between two codons, AATGAC, resulting in replacing Asn(AAT)Asp(GAC) to Asn(AAC). Another notable variant is a 382-bp deletion g.27848_28229del, almost completely removing ORF8, described by Su et al., 2020, supported by 14 samples from Singapore. Totally there are 4 indels that can be seen in 10 or more samples, 8 indels in 5 or more samples, 17 indels in 3 or more samples and 16 indels confirmed by just two samples. Some of these indels may affect protein functions - such as p.Val143_Tyr144delinsAsp and p.Tyr145del in spike protein and deletion of the first 7 amino acids in ORF10.
