## Supplemental Table 2 for "Quality control of low-frequency variants in SARS-CoV-2 genomes"

| Sequencing technology | Assembly method | # of genomes | # of genomes with singletons | # of genomes with >1 singletons |
| --- | --- | --- | --- | --- |
| All | - | 7862 | 19.49 | 5.47 |
| Illumina | - | 5602 | 18.58 | 5.23 |
| - | Phe pipeline | 1749 | 13.15 | 3.03 |
| - | Mapping | 1718 | 18.86 | 4.42 |
| - | Other/Not specified | 652 | 23.16 | 7.98 |
| - | Custom pipeline | 325 | 15.69 | 5.23 |
| - | Artic pipeline | 316 | 21.52 | 5.7 |
| - | iVAR | 281 | 17.08 | 2.49 |
| - | CLC | 258 | 34.11 | 15.12 |
| - | Seattle pipeline | 122 | 11.48 | 2.46 |
| - | MEGAHIT | 109 | 46.79 | 22.94 |
| - | SPAdes | 72 | 22.22 | 4.17 |
| Nanopore | - | 1609 | 19.14 | 4.79 |
| - | Other/Not specified | 837 | 20.07 | 4.3 |
| - | Artic pipeline | 509 | 16.5 | 4.52 |
| - | CLC | 184 | 25 | 7.61 |
| - | Custom | 77 | 12.99 | 5.19 |
| - | Mapping | 2 | 0 | 0 |
| Other/Not specified | - | 499 | 29.06 | 8.82 |
| - | Other/Not specified | 428 | 26.64 | 8.18 |
| - | CLC | 31 | 45.16 | 6.45 |
| - | Mapping | 22 | 36.36 | 18.18 |
| - | SPAdes | 11 | 36.36 | 18.18 |
| - | iVAR | 6 | 66.67 | 16.67 |
| - | MEGAHIT | 1 | 100 | 0 |
| Ion Torrent | - | 94 | 24.47 | 10.64 |
| - | Other/Not specified | 55 | 27.27 | 10.91 |

| | | | | |
|---|---|---|---|---|
| - | SPAdes | 21 | 4.76 | 0 |
| - | CLC | 12 | 41.67 | 25 |
| - | Custom | 4 | 25 | 25 |
| - | Mapping | 2 | 50 | 0 |
| Sanger | - | 42 | 26.19 | 11.9 |
| - | Other/Not specified | 42 | 26.19 | 11.9 |
| DNBSEQ | - | 16 | 25 | 6.25 |
| - | SPAdes | 11 | 18.18 | 9.09 |
| - | Mapping | 4 | 50 | 0 |
| - | Other/Not specified | 1 | 0 | 0 |

**Supplemental Table 2.** Full table with percentage of singleton-containing genomes depending on sequencing and assembly method. ”Other” category includes all custom pipelines, rarely used tools and samples with incomplete or absent information about assembly methods. “Mapping” category includes all assembly methods with words “mpileup”, “bwa”, “bowtie”, or “mapping” in the description.
